# Granulins modulate liquid-liquid phase separation and aggregation of TDP-43 C-terminal domain

**DOI:** 10.1101/812115

**Authors:** Anukool A. Bhopatkar, Vladimir N. Uversky, Vijayaraghavan Rangachari

## Abstract

Tar DNA binding protein (TDP-43) has emerged as a key player in many neurodegenerative pathologies including frontotemporal lobar degeneration (FTLD) and amyotropic lateral sclerosis (ALS). Important hallmarks of FTLD and ALS are the toxic cytoplasmic inclusions of C-terminal fragments of TDP-43 (TDP-43CTD), which are formed upon proteolytic cleavage of full-length TDP-43 in the nucleus and subsequent transport to the cytoplasm. TDP-43CTD is also known to form stress granules (SGs) by coacervating with RNA in cytoplasm under stress conditions and are believed to be involved in modulating the pathologies. Among other factors affecting these pathologies, the pleiotropic protein called progranulin (PGRN) has gained significant attention lately. The haploinsufficiency of PGRN, caused by autosomal dominant mutations in *GRN* gene, results in its loss-of-function linked to FTLD and ALS. But precisely how the protein contributes to the pathology remains unknown. Recently, cleavage to GRNs were observed to be a significant part of FTLD and ALS progression with specific GRNs exacerbating TDP-43-induced toxicity in *C.elegans*. In this report, we show that GRNs −3 and −5 directly interact with TDP-43CTD to modulate latter’s aggregation or stress granule formation in disparate ways in vitro. These results constitute the first observation of direct interaction between GRNs and TDP-43 and suggest a mechanism by which the loss of PGRN function could lead to FTLD and ALS.

## INTRODUCTION

Frontotemporal lobar degeneration (FTLD) is a neurodegenerative disorder characterized by progressive changes in behavior, speech, and personality among the elderly patients (1,2). These changes are bought upon by a gradual atrophy of the frontal and anterior temporal lobes of the brain (3–6). Based on the histopathology, heritable FTLD can be classified into four subtypes: FTLD-TDP form, which is characterized by the presence of cytoplasmic inclusions of TAR DNA binding protein-43 (TDP-43, also known as TARDBP), and which constitutes about half of all cases of FTLD (7); FTLD-tau in which cytoplasmic inclusions of hyperphosphorylated tau are observed (8–11); the third form of FTLD (FTLD-FUS) is associated with inclusions of fused in sarcoma (FUS) (8), while the last form is associated with ubiquitin positive inclusions (FTLD-U) (12). Inclusions of TDP-43 are also observed in patients with another neurodegenerative disorder called amyotropic lateral sclerosis (ALS) (13–15). Since TDP-43-based neuropathology is present in ~50% of FTLD and ~97% of ALS cases (16), TDP-43 represents a molecular link connecting these neurodegenerative diseases as a clinicopathological spectrum of the proteinopathies (14,17–19). It was also pointed out that TDP-43 may contribute to the pathogenesis of Alzheimer’s disease (AD) via both β-amyloid (Aβ)-dependent and Aβ-independent pathways (20). Furthermore, TDP-43 abnormalities have also been associated with traumatic brain injury (chronic traumatic encephalopathy) in both pre-clinical and clinical studies (7) and observed in cognitively impaired persons in advanced age with hippocampal sclerosis, Huntington’s disease and some other maladies (21). Because of the broad involvement of TDP-43 in pathogenesis of various neurodegenerative diseases, all such TDP-43-related abnormalities are grouped now into the TDP-43 proteinopathies (21,22).

TDP-43 is a member of the heterogenous ribonucleoprotein family involved in transcriptional regulation and mRNA splicing in neuronal and embryonic development (23). The protein consists of two RNA recognition motifs (RRMs) followed by a low-complexity, glycine-rich C-terminal domain (24). Intrinsic disorder predisposition of TDP-43 and four other proteins (superoxide dismutase 1 (SOD1), an RNA-binding protein fused in sarcoma (FUS), a cofilin-binding protein C9orf72, and polypeptides generated as a result of its intronic hexanucleotide expansions, and actin-binding profilin-1 (PFN1)), which are considered as the major drivers of ALS and FTLD pathogenesis, was recently evaluated (25,26). Based on this multifactorial analysis, it was concluded that the presence of significant levels of intrinsic disorder represents one of the common features to these proteins, and that intrinsic disorder plays a number of significant roles in their normal and pathological functions (25,26). Furthermore, the roles of intrinsic disorder of the TDP-43 and several other ALS- and FTLD-related proteins in the physiological and pathological liquid-liquid phase transitions (LLPTs) leading to the formation of various proteinaceous membrane-less organelles (PMLOs) was recently reviewed, indicating that intrinsic disorder and LLPTs are related to both normal and pathological functions of these proteins (26).

TDP-43 is predominantly localized in the nucleus, but is known to shuttle to the cytoplasm, where its precise roles remain unclear (24). It is however in the diseased state that the full-length protein is proteolytically cleaved generating 25 and 35 kDa (C25 and C35) C-terminal fragments that are transported into the cytoplasm, where they form inclusions observed in FTLD-TDP (27,28). The observed inclusions of TDP-43 C-terminal domain (TDP-43CTD) occur in the genetic backdrop of autosomal dominant, heterozygous mutations of *PGRN* that leads to haploinsuffiency of the protein. Homozygous *PGRN* mutations on the other hand lead to neuronal ceroid lipofuscinosis (NCL), a lysosomal storage disease (29). Under stress, TDP-43CTD is also known to bind RNA and get transported out of the nucleus to cytosol, where they undergo the phenomenon of liquid-liquid phase separation (LLPS) to form cytoplasmic granules as membraneless organelles (28). The process of LLPS has been identified in the past decade as a ubiquitous modulator of multiple cellular functions (30–39). Mechanistic details on the dynamics of such granules in TDP-43 pathophysiology remains unclear. However, many different proteins are known to interact with and partition into the phase separated droplets of TDP-43 (40,41). In FTLD and ALS, the aberrant LLPS of TDP-43 has emerged as a driving factor for the aggregation of the protein (40,42,43).

Progranulin (PGRN) is a 63.5 kDa secreted, glycosylated protein expressed in many cells including the neurons and in microglia (44–46), with pleiotropic roles in both physiological and pathological processes (47–51). The protein consists of seven and a half cysteine-rich domains called granulins (GRNs) (Fig 1a), (47,49,52–54) which are generated by proteolytic processing of PGRN by many proteases. PGRN and GRNs are observed to exist together *in vivo*, where they have opposing inflammatory functions (55). In FTLD, PGRN has been implicated in affecting the toxicity of TDP-43 by triggering its cleavage (56). Furthermore, the production of GRNs in haploinsufficient, but not the null state, suggests that they are key players in the disease phenotypes. The loss of PGRN function could arise from the increased proteolytic processing of the protein, which generates GRNs. Indeed, GRNs are shown to directly interact with TDP-43 and exacerbate latter’s levels and toxicity in *C.elegans*, establishing that PGRN cleavage to GRNs could represent an important part of the disease process in FTLD (57). Despite the growing evidence on the involvement of GRNs, their precise mechanisms in FTLD and related pathologies remain unclear. In this report, we sought to understand the interactions between GRNs and TDP-43CTD under reducing and oxidizing conditions *in vitro*, and whether the former is able to modulate the formation of liquid droplets or insoluble inclusions of TDP-43CTD, which shed insights into the role of GRNs in FTLD and ALS pathologies.

**Fig 1:**
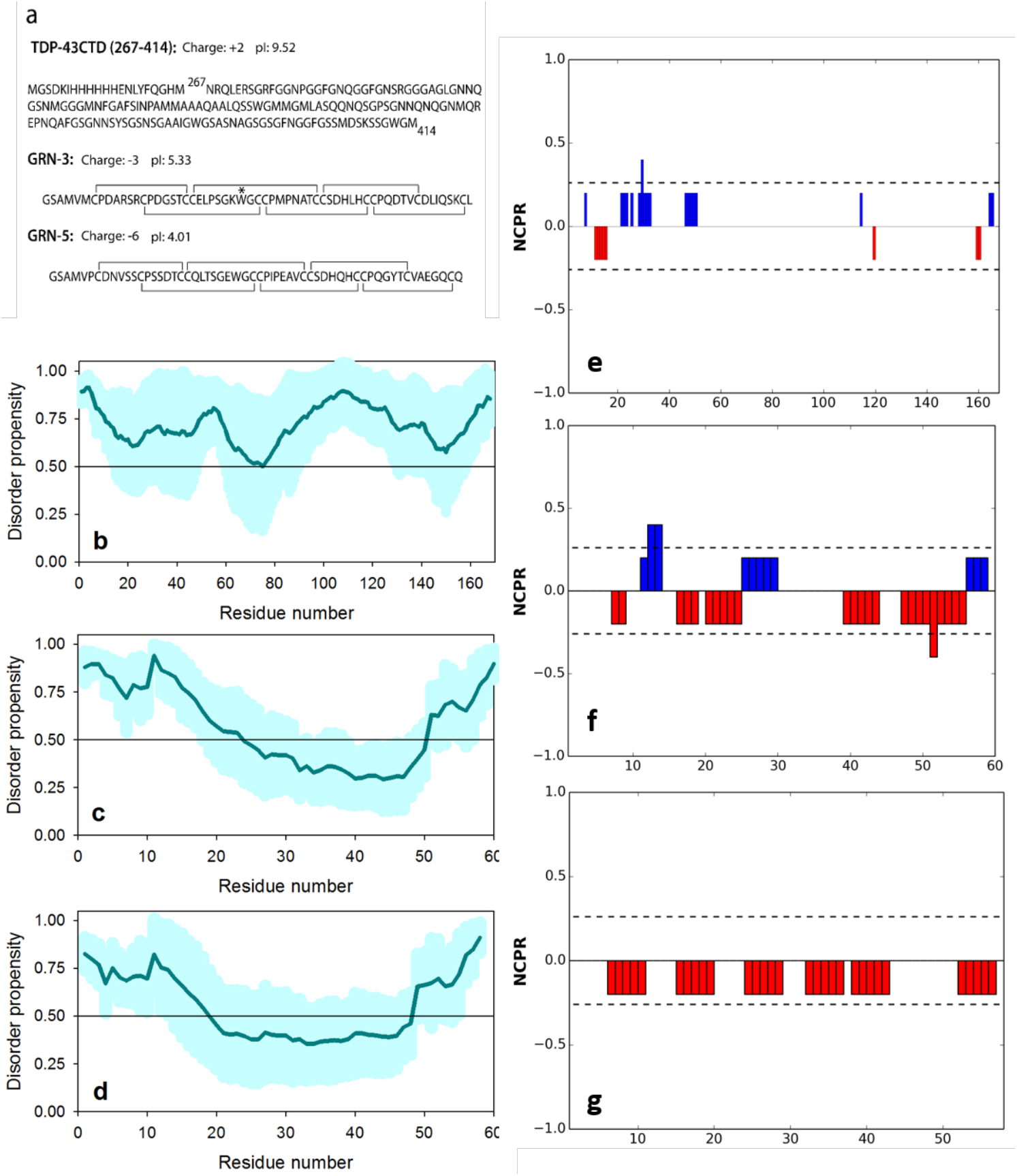
Sequence, disorder and charge distributions of the proteins used in this study. a) The sequence of TDP-43 C-terminal domain (TDP-43CTD) (267-414; 14.5 kDa) which constitutes the major part of the toxic 25 kDa TDP-43 C-terminal fragment with a hexa histidine-tag on the N-terminus. Sequences of GRN-3 and GRN-5 with putative disulfide bonds. A conservative mutation of Y to W (indicated as *) was made to GRN-3, which is a conserved residue in all GRNs. Both GRNs were used in fully reduced (free thiols) and oxidized conditions in this study. Computational evaluation of the per-residue intrinsic disorder predispositions of TDP-43CTD (b), GRN-3 (c), and GRN-5 (d). Here, the outputs of PONDR® VLXT, PONDR® FIT, PONDR® VL3, PONDR® VSL2, IUPred_S and IUPred_L were averaged to generate mean disorder profiles of different computational data query proteins. Diagrams of linear net charge per residue for TDP-43CTD (e), GRN-3 (f), and GRN-5 (g) generated by CIDER computational platform (http://pappulab.wustl.edu/CIDER/).

## RESULTS

### GRN-5 but not GRN-3 initiates LLPS of TDP-43CTD

The 14.5 kDa construct of TDP-43 that constitutes residues 267-414 in the C-terminal domain (TDP-43CTD) is a major part of the C25 proteolysis fragment that forms aberrant inclusions within the cytosol in FTLD and ALS patients (Fig 1a)(15). Fig 1b shows that TDP-43CTD is largely disordered, and previous study indicated that this protein contains sequence of low-complexity that is involved in LLPS (58). On the other hand, GRNs (1–7) are small ~ 6 kDa proteins with the conserved sequence; X_2-3_CX_5-6_CX_5_CCX_8_CCX_6_ CCXDXXHCCPX_4_CX_5-6_CX (Fig 1a). The 12 conserved cysteine residues form six putative intramolecular disulfide bonds (59) (Fig 1a). Fig 1c and Fig 1d show that despite the fact that these proteins contain high levels of cysteine, which is the strongest order-promoting residue, GRN-3 and GRN-5 are predicted to be characterized by high levels of intrinsic disorder. This is in line with the results of our previous studies, where we have shown that under reducing conditions, GRN-3 is fully disordered while in the oxidizing conditions, the protein obtains thermal stability via disulfide bonds without significant gain in the overall structure (60,61).

As a first step in characterizing the potential interactions between TDP-43CTD and GRNs, here we report those with GRNs 3 and 5 in both redox conditions (i.e., in completely reduced and oxidized states). GRN-5 was expressed and purified using the established method for GRN-3 (60) with some minor modifications. HPLC fractionation of GRN-5 resulted in multiple peaks (Fig S1a). Characterization of the major peak (arrow; Fig S1a) showed some degree of multimerization of GRN-5 detected in MALDI-TOF spectra perhaps by intermolecular disulfide bonds (Fig S1b). This was supported by Ellman’s and iodoacetamide analysis that showed up to two cysteines remained reduced (Fig S1c). Fractions 29-32 (Fig S1a) were pooled and treated with excess tris(2-carboxyethyl)phosphine (TCEP) to generate protein under fully reducing conditions (rGRN-5). The ^1^H-^15^N heteronuclear multiple quantum coherence (HMQC) NMR spectra of reduced rGRN-5 shows a narrow spectral dispersion between 7.5 and 8.6 ppm in ^1^H, and 108 and 130 ppm in ^15^N dimensions, indicative of absence of secondary structures and a disordered protein (62,63) (Fig S1d). HMQC spectra of the oxidized GRN-5 also show similar dispersion but loss of peaks was observed, which is likely due to the disorder and/or aggregation (Fig S1e). Both GRN-5 and rGRN-5 show a random coil secondary structure in far-UV CD (Fig S1f).

The primary sequence of GRN-5 is enriched in negatively charged residues without positively charged residues. As a result, the protein has a pI of 4.01. On the other hand, TDP-43CTD has an appreciable number of positively charged amino acids with a net pI of 9.52 (Fig 1a). Fig 1e and Fig 1g represent linear net charge per residue diagrams for these proteins generated by CIDER computational platform (64). Fig 1e shows that positively charged residues are preferentially concentrated within the 50 N-terminal residues of TDP-43CTD, whereas Fig 1g illustrates that negatively charged residues are distributed over the entire sequence of GRN-5. Importantly, ANCHOR analysis (65,66) of the TDP-43CTD construct utilized in this study revealed the existence of several disorder-based binding regions (residues 1-40, 66-99, 115-123, and 135-145). These regions represent molecular recognition features (MoRFs), which are intrinsically disordered segments that undergo folding at interaction with specific binding partners. Based on the counter-ionic character of the two proteins, we hypothesized that GRN-5 could potentiate interactions with TDP-43CTD to trigger LLPS. In addition, GRN-3 was chosen to test our hypothesis as this protein is also intrinsically disordered and has a pI of 5.33, but it contains both positive and negatively charged residues (Fig 1a and Fig 1f).

We also checked the amino acid sequence-based predispositions of TDP-43CTD, GRN-3, and GRN-5 to undergo biological LLPS. To this end, the CatGRANULE algorithm was utilized that predicts liquid-liquid phase separation propensity of a query protein based on the analysis of the phase separation features linked to its primary sequence composition, structural disorder, and nucleic acid binding propensities (67). This analysis indicated that the CatGRANULE scores for TDP-43CTD, GRN-3, and GRN-5 were 3.88, −3.06, and −2.83, respectively, indicating that only TDP-43CTD has an intrinsic propensity for granule formation associated to liquid demixing under the physiological conditions.

To see whether GRNs affect TDP-43CTD phase separation, the two proteins were co-incubated in 2:1 molar excess respectively and samples were monitored for turbidity increase and droplet formation by differential interference contrast (DIC) microscopy in both oxidizing (GRN-3 or GRN-5) and reducing (rGRN-3 or rGRN-5) conditions (Fig 2). At 37 °C, 20 μM TDP-43CTD buffered in 20 mM MES at pH 6.0 in presence of two molar excess of GRN-5 and rGRN-5 formed a turbid solution instantly, similar to the known phase separation shown by TDP-43CTD-RNA mixture (Fig 2a). On the other hand, incubation of TDP-43CTD by itself or with two molar excess of GRN-3 or GRN-3 did not show turbidity (Fig 2a). To visualize the phase separation and the presence of liquid droplets, the samples were observed under a DIC microscope. The TDP-43CTD control along with the co-incubated mixture containing GRN-3 or rGRN-3 did not show the formation of liquid droplets for up to 6 hours (Fig 2b). In contrast, liquid droplets were observed almost instantaneously when TDP-43CTD was co-incubated with two-fold molar excess of rGRN-5, GRN-5 or RNA (Fig 2b). The droplets that were numerous and small initially, grew in size by coalescing with one another in the next six hours of incubation, displaying fluid-like characteristics. To evaluate the effect of GRN concentration on TDP-43CTD phase separation, turbidity was measured within 10-15 minutes of incubation. Both GRN-5 and rGRN-5, show a concentration dependent increase in the turbidity, which saturates at two molar excess of GRN (Fig 2c). TDP-43CTD in presence of GRN-3 and rGRN-3 has no marked increase in turbidity even at high molar excess of GRNs (Fig 2c). As expected, droplets of TDP-43CTD and RNA showed high turbidity as compared to TDP-43 alone (Fig 2c-d). Co-incubations of two-molar excess GRN with TDP-43CTD indicated high turbidity for both GRN-5 and rGRN-5, with the former displaying the highest levels of turbidity (Fig 2d). Co-incubations of two-fold molar excess of GRN-5 or rGRN-5 with a mixture of RNA and TDP-43CTD showed an increase in turbidity as compared to TDP-43CTD-RNA control, with sample containing GRN-5 showing the highest effects (Fig 2d). Two molar excess of GRN-3 or rGRN-3 with TDP-43CTD showed no turbidity increase (Fig 2d). Similar addition of GRN-3 or rGRN-3 to TDP-43CTD-RNA mixture resulted in a slight decrease in turbidity levels as compared to the control (Fig 2d). Together, the data suggest GRN-5, and not GRN-3, is able to initiate or enhance the phase separation of TDP-43CTD alone or with RNA.

**Fig 2:**
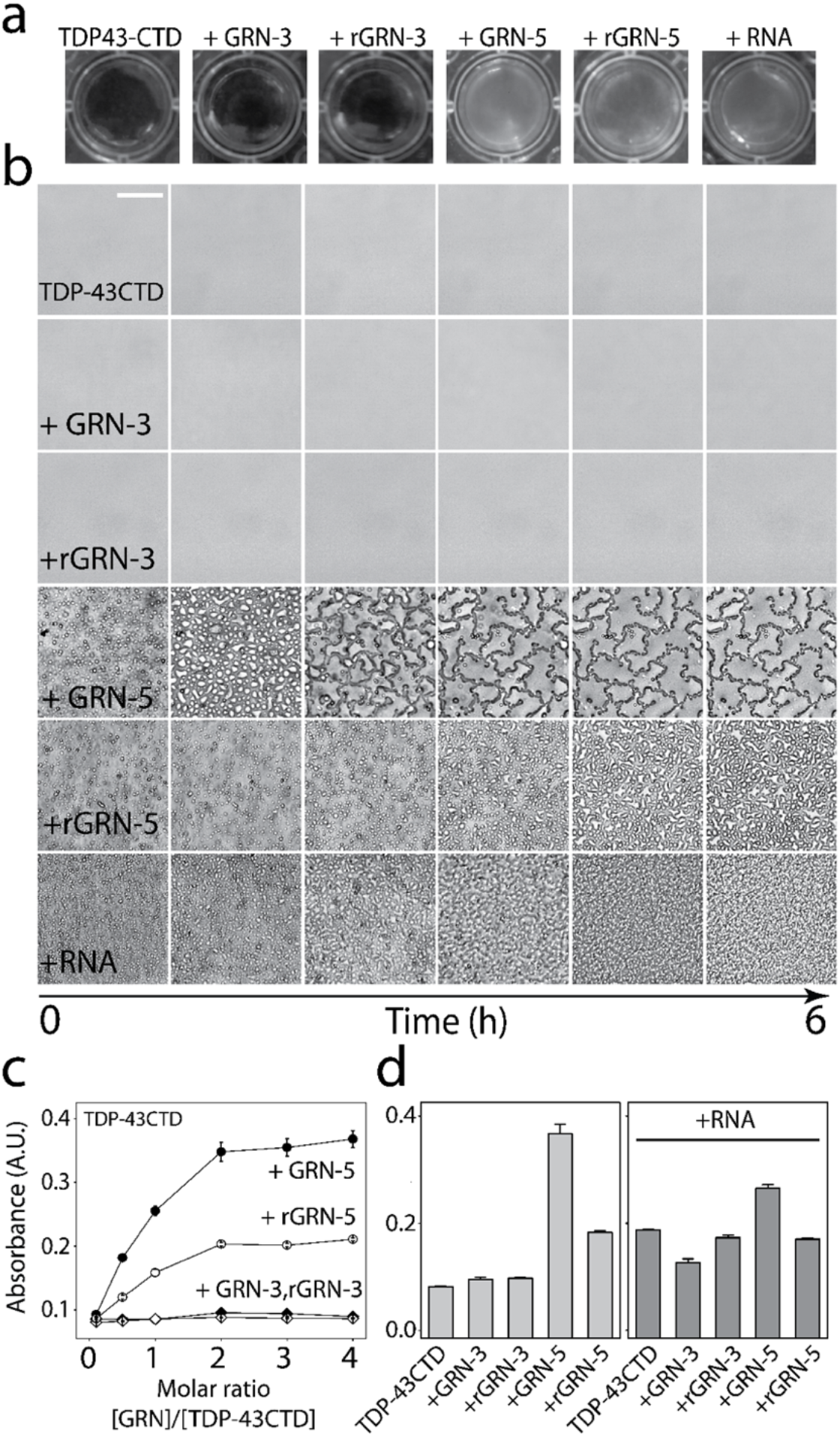
TDP-43CTD phase separates in the presence of both oxidized and reduced GRN-5. a) Visual snapshots immediately after the incubations of 20 μM TDP-43CTD in 20 mM MES buffer pH 6.0 by itself or with 40 μM granulins in oxidized (GRN-3; GRN-5) and reduced (rGRN-3; rGRN-5) conditions along with the positive control with 40 μg/mL RNA (stress granules). Only incubations with GRN-5, rGRN-5 and RNA show a turbid solution indicating phase separation. b) DIC microscopy images for the same reactions monitored for 6 hours. TDP-43CTD by itself or with GRN-3/rGRN-3 shows the absence of liquid droplets which those with GRN-5 or rGRN-5 coalescence of the droplets to varying degrees similar to the positive control (with RNA) confirming liquid-liquid phase separation. Scale bar represents 50 μm. c) Turbidity of the aforementioned reactions measured at 600 nm within ten minutes of incubation (n=3) and, d) Turbidity shown as bar graphs for the co-incubated solutions of 20 μM TDP-43CTD with 40 μM GRN-3 or 5 (reduced and oxidized), in the presence or absence of 40 μg/mL of RNA (n=3).

### GRNs −3 and −5 differently modulate the LLPS and formation of ThT-positive aggregates of TDP-43CTD

In FTLD and ALS pathologies, TDP-43CTD is known to form toxic inclusions in the cytosol. Yet, ambiguity remains regarding the nature of these inclusions with the presence of both thioflavin-T (ThT)-positive as well as ThT-negative aggregates reported (68,69) (70,71). To see how GRNs affected the formation of TDP-43CTD inclusions, we monitored ThT fluorescence for TDP-43CTD in the presence of increasing concentrations of GRN-3, rGRN-3, GRN-5, or rGRN-5 at 37 °C (Fig 3). Incubation of 20 μM TDP-43CTD alone in 20 mM MES buffer at pH 6.0 showed a typical sigmoidal increase in fluorescence with a lag time of ~ 8 hours (●; Fig 3) indicating the formation of ThT-positive species. For GRN-3, increasing stoichiometry from 0.1-molar excess (2 μM) to 4-molar excess (80 μM) showed increasing lag time for TDP-43CTD aggregation (Fig 3a); the lowest being 0.1x that showed a lag time of 12 hours, while the highest degree of increase was with 4-fold excess GRN-3, that showed a lag time of ~22 hours. Similarly, under fully reducing conditions, 0.1 to 4-fold excess of rGRN-3 also inhibited TDP-43CTD aggregation, but to a much lesser degree than GRN-3, with lag times ranging between 8 and 15 hours (Fig 3b). Similarly, incubation of GRN-5 in increasing concentrations (0.1 to 4-fold excess) with TDP-43CTD showed an overall inhibition of the TDP-43CTD fibrillation by increasing the lag times (12 to 36 hours, respectively) (Fig 3c). Incubations of rGRN-5 with TDP-43CTD, on the other hand, showed no discernable lag time but an instantaneous linear increase in ThT fluorescence for all stoichiometric incubations (Fig 3a; inset). These data suggest that GRN-3 and GRN-5 interact with TDP-43CTD differently to modulate latter’s aggregation. Furthermore, the observed LLPS of TDP-43CTD in the presence of GRN-5 and rGRN-5 (Fig 2) raises the possibility of the ThT dye partitioning into the droplets formed resulting in the higher ThT fluorescence observed at the outset. To investigate this, the reactions were visualized by fluorescence microscopy using another amyloid binding dye called thioflavin-S (ThS). Both ThT and ThS are known to detect amyloid aggregates (72,73). A 10 μM buffered solution of ThS was added to the co-incubated samples containing 20 μM TDP-43CTD and 40 μM GRNs; GRN-3 (Fig 3e), rGRN-3 (Fig 3f), GRN-5 (Fig 3g), and rGRN-5 (Fig 3h). Instantaneously upon addition, both GRN-3 and rGRN-3 showed no aggregate formation, and a disperse blue fluorescent haze was observed (Fig 3e and Fig 3f; 0h). The same reactions after 36 hours showed more concentrated areas where ThS was observed that coincided with the presence of insoluble aggregates (Fig 3e and Fig 3f; 36h). GRN-5 or rGRN-5 on the other hand, showed the presence of ThS within the droplets even at 0 hours (Fig 3g and Fig 3h; 0h), which continued to be present even after 36 hours (Fig 3g and Fig 3h; 36h). These data suggest that, *a*) ThS (or ThT) could partition into the droplets, and *b*) such partitioning results in the increase in fluorescent signals. In other words, the increase in ThT signals observed for both GRNs (Figs 3a-d) indicate different biophysical processes; while the presence of GRN-3 or rGRN-3 lead to insoluble aggregates, those of GRN-5 or rGRN-5 result in the droplet formation that could change to gelation or aggregates over time.

**Fig 3:**
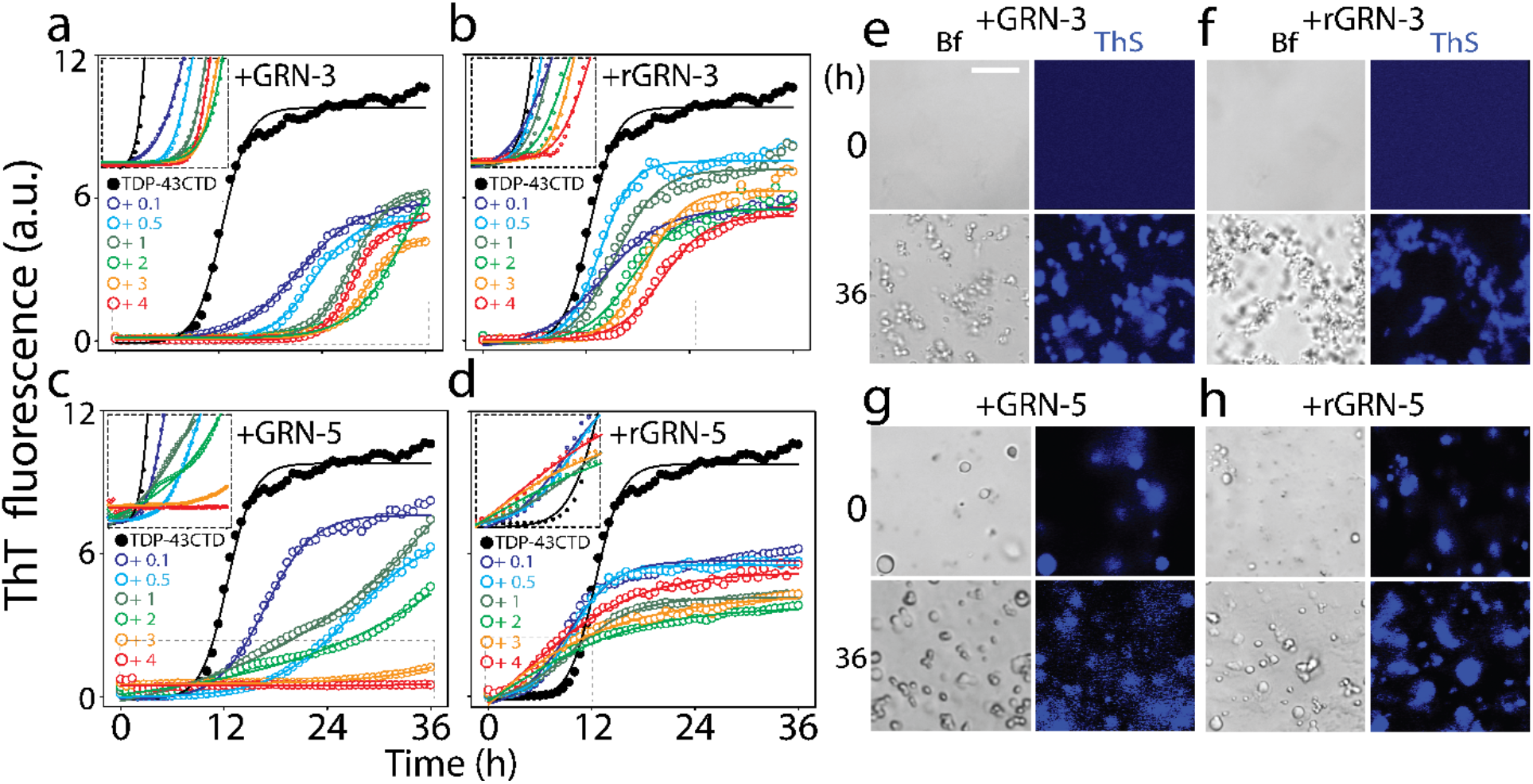
Modulation of TDP-43CTD aggregation by GRNs. a-d) The aggregation kinetics of 20μM TDP-43CTD in 20 mM MES buffer pH 6.5 at 37 °C with varying molar equivalence (2 to 80 μM corresponding to 0.1 to 4-fold molar excess) of GRN-3 (a), rGRN-3 (b), GRN-5 (c) and rGRN-5 (d), monitored by thioflavin-T (ThT) fluorescence. The inset shows the enlarged areas (boxed with dashed lines) highlighting the lag times during aggregation. The data was fitted to Boltzmann sigmoidal function. e-h) Localization of the aggregate-specific dye, thioflavin-S (ThS) observed by confocal fluorescence microscopy images on the two molar excess reaction samples from a-d immediately (0h) and after 36 hours of incubation. Scale bar represents 20 μm and brightfield view is denoted by Bf.

### Both GRNs colocalize with TDP-43CTD within the droplets or aggregates

To further probe whether the droplets are complex coacervates of TDP-43CTD and GRNs or formed without GRNs’ partitioning into the separated phase, fluorescence microscopy was used to observe the colocalization of the two proteins. TDP-43CTD and GRNs, labeled with HiLyte 647 and HiLyte 405, respectively, were incubated separately under oxidizing and reducing condition and monitored for 36 hours (Fig 4). As expected, TDP-43CTD by itself did not form phase separated droplets immediately after incubation (Fig 4a), and over the next 36 hours, sparse deposits of aggregated protein were observed (Fig 4a). Co-incubations of GRN-3 or rGRN-3 with TDP-43CTD showed no LLPS immediately after incubation (0h; Fig 4b and Fig 4c), parallel to the observation by DIC and turbidity (Fig 2). Even after 36 hours, no phase separated liquid droplets, but insoluble, fibrillar deposits were observed (18 and 36h; Fig 4b and Fig 4c) that bind amyloid dyes as shown above (Fig 3). Moreover, the GRNs were observed to be localized in the vicinity of TDP-43CTD deposits as well as within them (36 h; Fig 4b and Fig 4c). On the other hand, co-incubations of both GRN-5 and rGRN-5 with TDP-43CTD resulted in the emergence of well-defined liquid droplets immediately after incubation (0h; Fig 4d and Fig 4e). The number of droplets increased during the next 36 hours of incubation, with some becoming more distorted spherical droplets (18 and 36 h; Fig 4d and Fig 4e). Importantly, both GRNs (blue) colocalize within the droplets formed by TDP-43CTD (red). To further glean into the physical nature of the droplets or insoluble inclusions formed, fluorescence recovery after photobleaching (FRAP) was conducted (Fig 4f-j). After 36h of incubation, TDP-43CTD control showed no recovery after bleaching indicative of insoluble solid nature of the aggregates (Fig 4f). A similar behavior was observed with the coincubations of TD-43CTD with GRN-3 or rGRN-3 (Fig 4g and 4h) suggesting that GRN-3 in both reducing and oxidized states promoted fibrils of TDP-43CTD as seen from the microscopy images. However, GRN-5 in both redox states (GRN-5 and rGRN-5) showed exponential fluorescence recovery immediately after incubation indicating fluid property of the droplet (**0h**; Fig 4i and 4j). After 36h, the extent of recovery was slightly mitigated indicating a possible “gelation” of the droplet (**36h**; Fig 4i and 4j). Together, the data further confirm that both GRN-5 and rGRN-5 induce LLPS with TDP-43CTD by coacervation into the droplets. On the other hand, GRN-3 does not induce LLPS but forms insoluble deposits of aggregated forms of TDP-43CTD, indicating the differential specificity for the two GRNs in interacting with TDP-43CTD.

**Fig 4:**
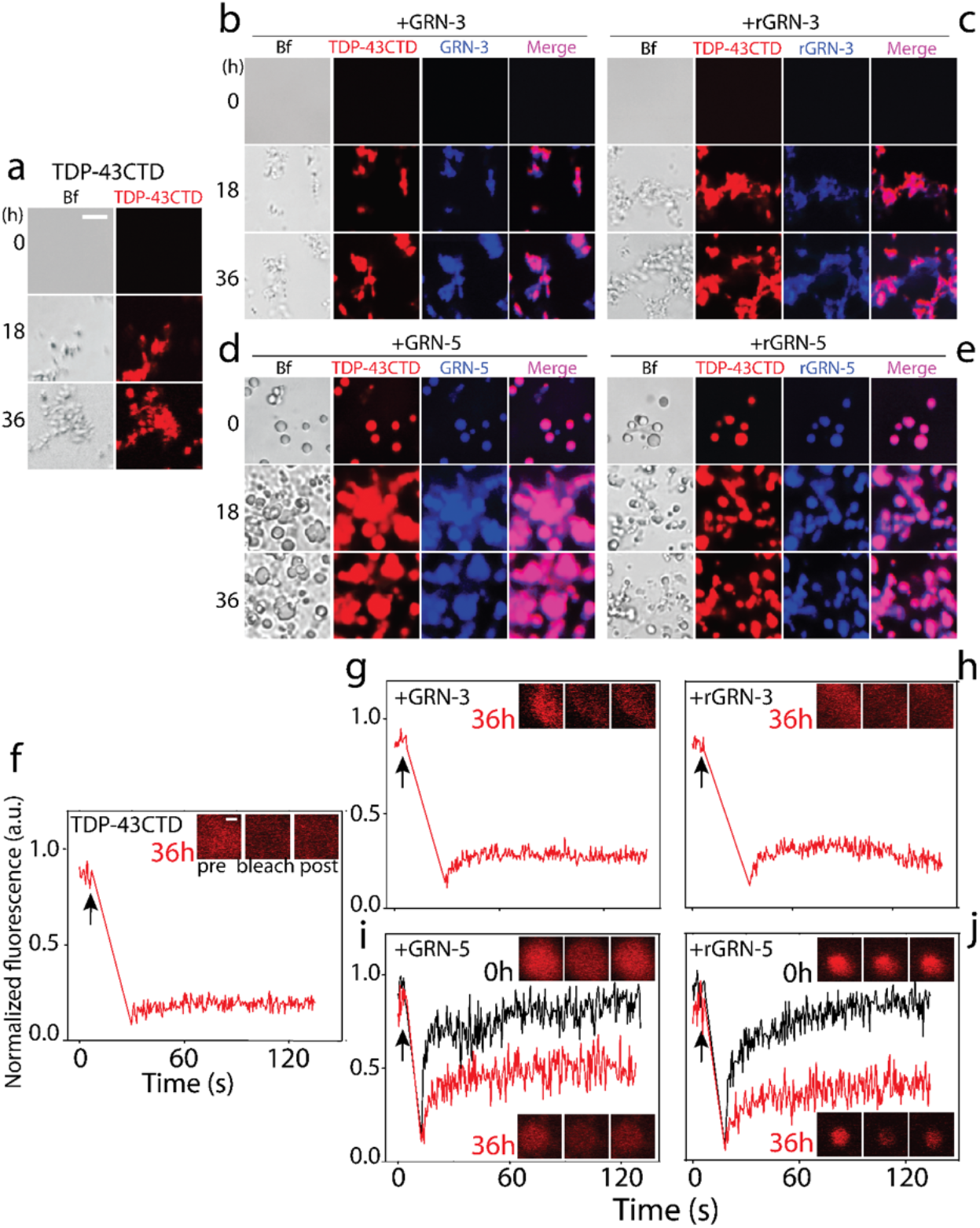
Coacervation of GRNs with TDP-43CTD and dynamics of phase separated droplets. Localization of TDP-43CTD and GRNs was monitored on a co-incubated mixture of the two proteins in 1:2 molar ratio. A 0.2 μM aliquot (1%) of TDP-43CTD labeled with HiLyte 647 fluorescent tag was mixed with 20 μM unlabeled TDP-43CTD in 20 mM MES buffer pH 6.5 at 37 °C and was co-incubated separately with a mixture of 40 μM GRNs containing 1 % of respective proteins labeled with HiLyte 405. The co-incubated mixture was visualized using DIC microscope. ‘Bf’ represents bright field in all the images. a) TDP-43CTD control in the absence of GRNs shows formation of fibrillar structures after a period of 18 hours that increase in size after 36 hours. TDP-43CTD co-incubated with GRN-3 (b) or rGRN-3 (c) show colocalization within insoluble fibrils over time with some peripheral localization also observed. Co-incubated samples containing GRN-5 (b) or rGRN-5 (c) show co-localization with TDP-43CTD in phase-separated liquid droplets. Scale bar represents 20 μm. f-j) Fluorescence recovery after bleaching (FRAP) data performed using HiLyte 647 labeled-TDP-43CTD. Arrow represents the time of bleaching. f) Aggregates formed by TDP-43CTD control show no fluorescence recovery after 36 hours. g-h) fluorescence recovery in presence of GRN-3(g) or rGRN-3(h) immediately (0h; black line) and after 36 h of incubation (red line). i-j) fluorescence recovery in presence of GRN-5 (i) or rGRN-5 (j) immediately (0h; black line) and after 36 h of incubation (red line). The confocal images in the insets in f-j show the area bleached before, after, and after recovery. Scale bar represents 2 μm.

To further confirm and quantify the partitioning and co-localization of TDP-43CTD and GRNs, the relative amounts of the proteins within the droplets as well as the deposits were quantified by MALDI-TOF mass spectrometry. Using a known amount of insulin (6.45 ng/μL) as an external standard, the mass spectra were obtained (Figs 5a-d) and the relative quantities of the proteins were calculated and normalized (detailed in Materials and Methods). Aliquots of the samples used in Fig 3 and Fig 4 were obtained after 36 hours for quantitative analysis. Both GRN-3 and rGRN-3 showed positive correlation between the amount of GRN co-incubated and TDP-43CTD with the amount of GRN-3 or rGRN-3 increasing in the sedimented pellet with increase in stoichiometry (Fig 5e and Fig 5f). More importantly, the ratio of GRN to TDP-43CTD remained below one, suggesting GRN-3 or rGRN-3 may augment TDP-43CTD aggregation by nucleation and not by forming a co-complex in stochiometric proportions. This observation is similar to the recently observed ability of GRN-3 and rGRN-3 to form fibrils of amyloid-β(1–42) (Aβ42) peptide involved in Alzheimer disease (AD), where minimal GRN was observed in the pellet (74). On the other hand, the ratio of GRN-to-TDP-43CTD obtained from the sedimented droplets of TDP-43CTD and GRN-5 or rGRN-5 co-incubations did not show a linear increase with increasing concentrations of GRN as observed with GRN-3 (Fig 5c and Fig 5d). In co-incubations of GRN-5 and rGRN-5, the GRN-to-TDP-43CTD ratio remained at 0.75 and 1.1, respectively, over the entire stoichiometric range. This indicates that a 1:1 complex is likely formed between GRN-5 /rGRN-5 and TDP-43CTD, which stabilize the droplets. It is known that the phenomenon of coacervation is dependent on a defined valency of interactions among the molecules which remain fixed within the dense phase (75–78). The data indicates that a 1:1 stoichiometry of GRN-5: TDP-43CTD could be sufficient to fulfil the required valency of the weak interactions that mediate the LLPS.

**Fig 5:**
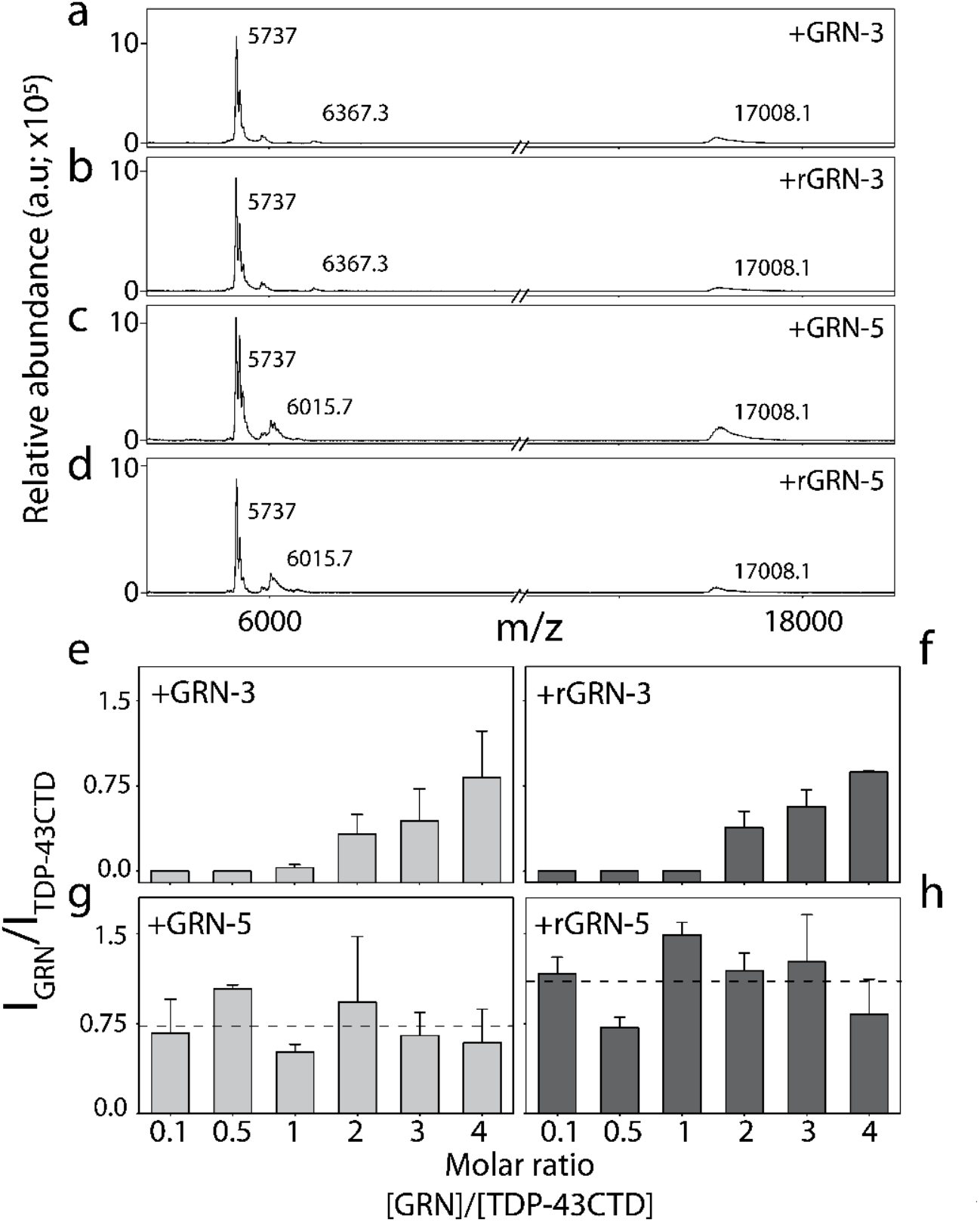
Quantitation of sedimented droplets and pellets from the co-incubations of TDP-43CTD and GRNs. The co-incubated samples of TDP-43CTD and GRNs from figures 3 and 4 were sedimented at 18,0000xg for 20 minutes after 36 hours of incubation, and the pellets were quantified by MALDI-Tof mass spectrometry using insulin (6.45 ng/μL, m.w 5737 Da) as an external standard. The amounts of GRN-3 (mw 6367.3 Da), GRN-5 (mw 6015.7 Da) or TDP-43CTD (mw 17008.1 Da) within the pellets were determined with respect to the standard (detailed in Materials and Methods) and plotted as a ratio of GRN to TDP-43CTD for the respective molar ratios. a-d) MALDI-ToF spectra of samples containing 20 uM TDP-43 with two molar excess of GRN-3 (a), rGRN-3 (b), GRN-5 (c) and rGRN-5 (d). e-h) Ratio of intensity of GRNs present in complex with TDP-43CTD in samples of varying molar ratios of the former shows an increase of GRN-3 (e) and rGRN-3 (f) within the complexes at higher molar excess, while GRN-5 (g) and rGRN-5 (h) are present in a constant amount. GRN-5 forms a 1:1 complex with TDP-43CTD in both redox forms; GRN-5 (0.73) and rGRN-5 (1.1) as indicated by the average amount (dashed line).

### GRNs modulate stress granules formed by TDP-43CTD and RNA

One of the main aspects of TDP-43 pathobiology is the coacervation with RNA molecules to form membraneless organelles in the form of stress granules (SG) (79–81). To see whether GRNs 3 and 5 are able to interact with, and modulate SG, torula yeast RNA extract was co-incubated with TDP-43CTD in the presence or absence of 1:2 molar excess of GRNs. As expected, the co-incubation of RNA with TDP-43CTD did not show any ThT increase as they form SG (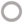; Fig 6a and Fig 6b) as compared to the TDP-43CTD alone (● Fig 6a and Fig 6b). Inclusion of rGRN-3 to the RNA and TDP-43CTD mixture significantly increased the rate of ThT-positive aggregates of DP-43CTD (Fig 6a) but slightly delayed the aggregation compared to TDP-43CTD alone or with rGRN-3 (Fig 6a). Similar augmentation of aggregation was observed with the co-incubation of GRN-3 and TDP-43CTD-RNA mixture (Fig 6a) although the increase in rate of aggregation was less pronounced with GRN-3 as compared to rGRN-3 as previously observed in Fig 3. Co-incubation of rGRN-5 to the TDP-43CTD-RNA mixture ‘rescued’ the aggregation of TDP-43CTD with a linear increase in ThT fluorescence observed within hours of incubation (Fig 6b). However, augmentation of ThT-positive aggregates was also pronounced with the co-incubation of GRN-5 (; Fig 6b). These data indicate that GRNs are able to modify SG to promote aggregates of TDP-43CTD. To further understand the mechanism of GRNs’ influence on SG, co-incubations of fluorescently-tagged samples were monitored by DIC microscopy. As expected, TDP-43CTD-RNA showed SG formation immediately after incubation that remained for 36 hours (Fig 6c). Addition of rGRN-5 to TDP-43CTD-RNA mixture also showed the formation of droplets immediately, which fused and grew bigger in size during the next 36 hours, with both TDP-43CTD and rGRN-5 colocalizing within the SG (Fig 6d). Addition of GRN-5 to TDP-43CTD-RNA mixture also led to droplet formation which grew in number but not in size during the same time (Fig 6e). GRN-5 also co-localized with TDP-43CTD within the SGs. Incubation of rGRN-3 and GRN-3 with TDP-43CTD-RNA mixture showed distortions in the morphology of SG accompanied with an overall reduction in the size during the course of the experiment (Fig 6f).

**Fig 6:**
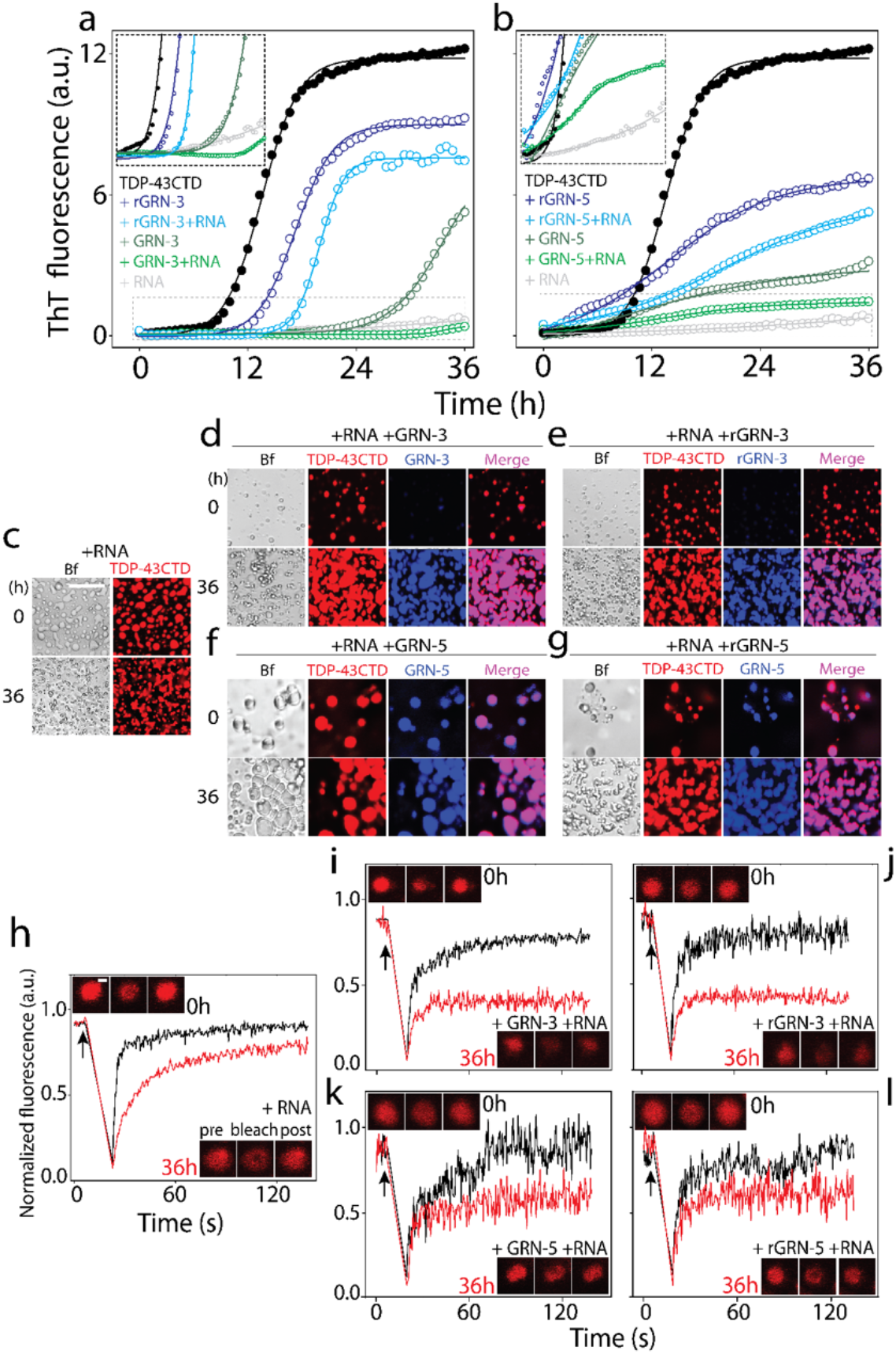
Modulation of TDP-43CTD-RNA stress granules by GRNs. **a-b)** Aggregation kinetics of 20μM TDP-43CTD in 20 mM MES buffer pH 6.5 at 37 °C with 40 μM of GRN-3 (a) and GRN-5 (b) in both oxidized and reduced forms, and in presence or absence of RNA (40 μg/mL) monitored by ThT fluorescence. The inset shows the enlarged areas (boxed with dashed lines) highlighting the lag times during aggregation. c-g) Localization of the protein from the reactions similar to those in panels a and b visualized by labeling TDP-43CTD and GRNs with HiLyte 647 and HiLyte 405, respectively, and in the presence and absence of unlabeled RNA by DIC microscopy. Oxidized (f) and reduced (g) forms of GRN-5 colocalize with TDP-43 into liquid droplets in presence of RNA. GRN-3 (d) and rGRN-3 (e) do not localize to phase separated droplets with TDP-43 even in presence of RNA but form solid complexes. FRAP analysis performed using HiLyte 647 labeled-TDP-43CTD with RNA and GRNs at initial timepoint (black line) and after a period of 36 hours (red line). Arrow represents the time of bleaching. h) Control SGs formed by TDP-43CTD and RNA. i-l) fluorescence recovery in presence of GRN-3(i), rGRN-3(j), GRN-5 (k) or rGRN-5 (l). The confocal images in the insets in h-l show the area bleached before, after, and after recovery. Scale bar represents 2 μm.

**Fig 7:**
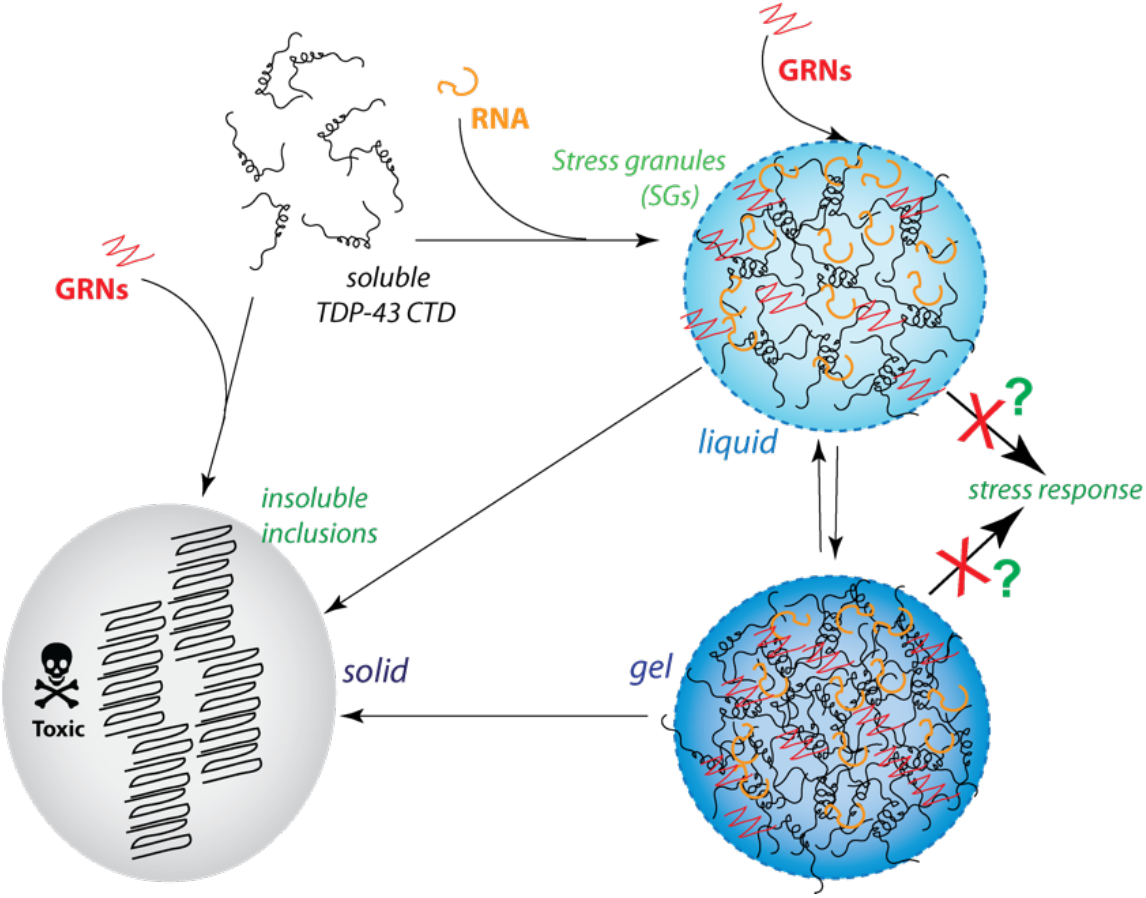
Schematic of the data presented in this study. GRNs selectively modulate the dynamics of TDP-43 aggregation or phase separation, which may have physiological implications. The question marks indicate our hypothesis and is not a result that is supported by this report.

To probe the internal dynamics of the SG droplets, FRAP was monitored temporally on labelled TDP-43CTD in presence of RNA and GRNs. The fluorescence recovery rates for the control SGs formed by TDP-43CTD and RNA showed near identical rates at both 0 and 36 hours, indicating preservation of internal mobility and dynamism over the incubation period (Fig 6h). However, SGs formed in presence of both redox forms of GRN-5 displayed a decreased fluorescence recovery after 36 hours indicating an inhibition of internal mobility, which could indicate the maturation of the droplets into gels, process commonly known as ‘gelation’ (Fig 6k; GRN5 and Fig 6l; rGRN-5). It is possible that these ‘gelated’ SG eventually form insoluble aggregates of TDP-43CTD, but we did not observe such aggregates within the window up to 48h of incubation (data not shown). The increase in ThT fluorescence (Fig 6b) can be explained by the fact that ThS (and likely ThT) partitions into the droplets and fluoresces (Fig 3). From these experiments it is clear that both GRNs 3 and 5 in redox conditions induce the formation of ThT-positive inclusions of TDP-43CTD but adopt two different mechanisms. GRN-5 seems to be enhancing the LLPS of the SGs leading gelation while GRN-3 disrupts it.

## DISCUSSION

Genetic conjunction of PGRN with FTLD/ALS is well-established with the implication of autosomal dominant mutations in PGRN leading to its haploinsuffiency. The loss of PGRN function is also speculated to arise at a posttranscriptional level (82) such as increased PGRN proteolysis. However, the role of GRNs in these pathologies remains unknown, despite their localization in FTLD and ALS brains. GRN immunopositivities were observed in a region-specific manner in both AD and FTLD human brains (83). In one report, GRNs were found in both microglia and neurons, but their distribution in the brain regions were asymmetrical (83). In FTLD, the production of GRNs in haploinsufficient and not null state suggests that they may be key players in the disease phenotypes. GRNs were shown to directly interact with TDP-43 and exacerbate latter’s levels and toxicity in *C.elegans*, establishing that PGRN cleavage to GRNs is an important part of the disease process in FTLD (57). Moreover, the recent discovery of PGRN/GRNs being associated with lysosomal dysfunction (84) posits the question whether GRNs could be involved in autophagic fate of TDP-43 inclusions under redox stress. The results presented here attempts to answer some of the key questions regarding GRNs and TDP-43CTD such as how GRNs modulate the dynamics between SG and fibril formation of TDP-43CTD and do SGs facilitate the formation of insoluble TDP-43CTD inclusions.

The findings of this study showcase how GRNs −3 and −5 modulate the dynamics of TDP-43CTD in forming insoluble aggregates or SGs in both oxidizing and reducing conditions. It is clear from the data that GRN-3 and GRN-5 affect TDP-43CTD aggregation via disparate mechanisms; while GRN-3 induces the formation of insoluble inclusions, GRN-5 coacervates with TDP-43CTD to undergo LLPS in both reduced and oxidized forms an interaction which is likely to be driven by the counter-ionic electrostatic characteristics of TDP-43CTD and the negatively charged GRN-5 which are known to be common mediators of LLPS (85–87). Furthermore, GRN-3 and GRN-5 also disrupt SGs formed by RNA and TDP-43CTD. Specifically, GRN-3 in both redox states accelerate the formation of ThT-positive, solid aggregates of TDP-43CTD. This observation supports the one by Salazar and co-workers who observed exacerbation of TDP-43 toxicity in *C.elegans* by GRN-3 and GRN-7 (57). GRN-5, on the other hand seems to remodify the SGs by inducing gelation, which eventually shows ThT-positive species. At this point, it is unclear as to what cellular ramifications such remodeling of SGs by GRN-5 could be and our on-going investigations will glean into this in the future. Nevertheless, our results provide the first in vitro evidence for a direct interaction between GRN and TDP-43CTD.

The increasing relevance of GRNs in FTLD and associated neurodegenerative disorders imparts significance to the results presented in this study. Despite its significance, how haploinsufficiency of PGRN results in TDP-43 inclusions in FTLD and ALS patients remains unknown. Common mutations associated with PGRN haploinsufficiency have been linked to mutant mRNA degradation (88) which several groups have identified leads directly to reduced circulating levels of PGRN (89–91). A few have also argued that increased proteolysis of PGRN as a reason for haploinsufficiency, which will directly correlate with increased GRN levels (57). A more recent study established that more effective proteolytic processing of cathepsin D, an aspartyl protease that is known to cleave PGRN, occurs in the presence of GRNs (92), which seem to support the idea that generation of GRNs to be the reason for decreased availability of PGRN. Results presented in this study establish a potential mechanism by which GRNs directly interact with TDP-43CTD and modulate the latter’s ability to aggregate and/or phase separate (Fig 6). It can be hypothesized that the PGRN proteolytic products directly but selectively interact with free TDP-43CTD to augment its aggregation towards cytotoxicity. Alternatively, under redox stress, GRNs can interact with SGs to modulate the dynamics within the droplets towards the formation of insoluble inclusions or lead to a decreased stress response. Whether and how these processes are regulated by phosphorylation and other mutations within GRNs and TDP-43 and are currently under investigation and will be reported at a later time.

## EXPERIMENTAL PROCEDURES

### Expression and purification of recombinant proteins

#### Granulins (GRN-3 & GRN-5)

Unlabeled GRN-3 was expressed and purified from *E. coli* SHuffle^TM^ cells (New England Biolabs) as described previously (61), while unlabeled GRN-5 and ^15^N labeled GRN-5 were expressed in Origami 2 DE3 (Invitrogen). Briefly, the proteins were expressed as a GRN:trxA fusion construct and purified using immobilized-nickel affinity chromatography. The fusion protein was then cleaved by addition of restriction grade thrombin (Bovine, BioPharm Laboratories) at 1U per 1 mg of the protein to remove both trxA and the His-tag. The reaction was incubated at room temperature (~25°C) for 22-24 hours. The protein was then fractionated on a semi-preparative Jupiter® 5 µm 10×250 mm C18 reverse phase HPLC column (Phenomenex), using a gradient elution of 60 – 80 % acetonitrile containing 0.1% TFA as previously described (61). The concentration of the proteins was estimated spectrophotometrically using molar extinction coefficients of 6250 M^−1^cm^−1^ for GRN-3 and 7740 M^−1^cm^−1^ for GRN-5 at 280 nm. The number of free cysteines was calculated from Elman’s assay and by iodoacetamide labeling as previously reported (61). rGRN-3 and rGRN-5 were generated by the addition of 2 mM Tris(2-carboxyethyl)phosphine hydrochloride (TCEP) to the HPLC fractionated protein for 2 hours at room temperature and were used as the reduced form. ^15^N labeled GRN-3,5 were generated by growing the cells in M9 minimal media enriched with ^15^NH_4_Cl.

#### TDP-43CTD

The plasmid for the TDP-43CTD expression was a gift from Nicolas Fawzi (Addgene plasmid # 98669; http://n2t.net/addgene:98669; RRID:Addgene 98669). The protein is expressed as a fusion construct with a His-tag (His_6_) at the N-Terminal followed by a TEV cleavage site The plasmid was expressed in *E.coli* BL21 Star™ (DE3) cells (Life Technologies). Transformed cells were grown at 37°C in LB medium supplemented with 100µg/ml of kanamycin. Overexpression of the protein was induced by adding isopropyl β-D-1-thiogalactopyranoside (IPTG) to a final concentration of 0.5 mM at an optical density (O.D_600_) of 0.5-0.7 AU. After overnight induction at room temperature (~25°C), cells were harvested by centrifugation (15000xg, 4 °C) and used immediately or stored at −20°C. The cells were resuspended in lysis buffer (20 mM Tris pH 8.0, 500 mM NaCl, 5 mM imidazole, 6 M urea). 0.5 mM phenylmethylsulfonyl fluoride (PMSF) was also added to the resuspension. Cells were lysed using Misonix XL-2000. The lysate was centrifuged at 20000 xg, 4°C, for 1 hour to remove cellular debris. The supernatant was incubated with Ni-NTA resin at 4°C for 2 hours. The slurry was resuspended in Kimble Kontes Flex column and washed with wash buffers (20 mM Tris pH 8.0, 500 mM NaCl, 6 M Urea) of 15 mM and 30 mM imidazole and protein was eluted in elution buffer (20 mM Tris pH 8.0, 500 mM NaCl, 6 M Urea, 150 mM imidazole). Eluate was buffer exchanged into storage buffer (20 mM Tris pH 8.0, 500 mM NaCl, 2 M Urea) and concentrated using Amicon Ultra-Centrifugal units (Millipore). Concentrated protein was flash frozen and stored at −80°C. Concentrated aliquots were thawed on ice and desalted into 20 mM MES pH 6.0 using Zeba Desalting Spin Columns (Thermo) and used as such for studies.

### Thioflavin-T fluorescence

The aggregation kinetics of TDP-43CTD were monitored using Thioflavin-T (ThT) on a BioTek Synergy H1 microplate reader. 20 µM TDP-43CTD under varying reaction conditions was incubated at 37°C for 36 hours in presence of 10 µM ThT. Samples were excited at 440 nm and emission was monitored at 480 nm. The ThT kinetic data was fit using the following Boltzmann function.

### MALDI-ToF mass spectrometry

Pellet and droplet characterization of the co-incubated protein complexes was performed on a Bruker Datonics Microflex LT/SH ToF-MS system. Samples from aggregation kinetic reactions were recovered after a period of 36 hours. The recovered samples were spun down at 18000 xg and supernatant was decanted. The pellet obtained was then resuspended in equal volume of reaction buffer (20 mM MES, pH 6.0). The pellet and supernatant fractions were then prepared for MALDI-MS by mixing with 6.5 ng of insulin (external standard). Additionally, the pellet samples were dissolved with formic acid in a 1:1 ratio to allow disaggregation of fibrils and other insoluble complexes. Both aliquots were then spotted onto a Bruker MSP 96 MicroScout Target with 1:1 ratio of sample:sinapinic acid matrix in saturated acetonitrile and water. Instrument calibration was performed using Bruker Protein Calibration Standard I (Bruker Daltonics).

### Differential Interference Contrast (DIC)

The phase separation was monitored using DIC on a Leica DMIL LED microscope under 20x or 60x magnification with a DIC polarizing filter. The reactions were observed on a clear-bottom Nunc™ MicroWell™ 96-Well Microplates (Thermo).

### Fluorescence microscopy

For colocalization studies of GRNs and TDP-43CTD during the course of their interactions, labeling of the respective proteins with amine-reactive fluorescent dyes was performed and visualization was done on a Zeiss LSM 510 Meta confocal microscope at 40x magnification under oil-immersion with a DIC polarizing filter. Briefly, TDP-43CTD was labelled with HiLyte-647 (AnaSpec) and GRN-3,5 were labelled with HiLyte-405 (AnaSpec). Free dyes were removed via the use of desalting columns; Zeba Desalting Spin Columns (Thermo) for labeled TDP-43CTD and Clarion™ MINI Spin Columns, Desalt S-25 (SorbTech) for labeled GRN-3,5. 20 uM TDP-43CTD was mixed with 40 µM GRN-3,5 or 40 ug/mL of RNA. A ratio of 1% Labeled: 99% Unlabeled proteins was used for both TDP-43CTD and GRNs. Reactions were performed on the following experimental setup: A glass cover-slip was attached onto one end of a two-side open ended 96 well plate (Greiner Bio-One). Reactions were loaded into a well and covered with an optically clear plate-sealing film (Thermo). Reactions were then monitored at certain timepoints. Similarly, for observing the localization of ThS, 10 µM ThS (AAT bioquest) was added to sample containing 20 µM TDP-43CTD with 2 molar excess of GRNs. Samples were visualized using the setup described above on a Zeiss LSM 510 Meta.

### FRAP

FRAP assays were performed in open-ended 96 well plates as described above; 10 µM TDP-43CTD was mixed with 20 µM GRN-3,5 or 20 µg/mL of RNA. Reactions were monitored 10 mins after mixing to allow deposition of suitable droplets onto the bottom of the well, which were used for bleaching. TDP-43CTD labelled with HiLyte-647 (AnaSpec) was used as the probe in the assays. LSM 510 Meta confocal microscope at 63x magnification under oil-immersion with a DIC polarizing filter and FRAP module. The region of interest was bleached with 633 nm laser line with 100% bleaching intensity for 750 iterations and imaging was performed at 5% laser intensity for 120 secs after bleaching. Numerical aperture was kept at 0.69 AU.

### NMR spectroscopy

The HMQC NMR spectra for 20 µM ^15^N GRN-5 or rGRN-5 resuspended in 20 mM MES (pH 6.0) with 10% D_2_O was acquired on a Bruker Advance – III-HD 850 MHz NMR spectrometer equipped with a Bruker TCI cryoprobe at the high field NMR facility of University of Alabama, Birmingham as described previously(61). For the HMQC NMR spectra of ^15^N GRN-5 or rGRN-5 in presence of TDP-43CTD, 20 µM GRN reactions were prepared with 0.5 mM TCEP (for rGRN-5), or without (for GRN-5) and latter was added to a final concentration of 10 µM immediately before acquiring the spectra.

### Computational analysis of TDP-43CTD, GRN-3, and GRN-5

Per-residue intrinsic disorder predispositions of TDP-43CTD, GRN-3, and GRN-5 were evaluated by a set of commonly used disorder predictors, such as PONDR^®^ VLXT(93), PONDR^®^ VSL2 (94), and PONDR^®^ VL3 (95) available on the PONDR site (http://www.pondr.com) PONDR^®^ FIT (96) accessible at the DisProt site (http://original.disprot.org/metapredictor.php), and the IUPred computational platform available at IUPred2A site (https://iupred2a.elte.hu/) that allows identification either short or long regions of intrinsic disorder, IUPred-L and IUPred-S (97–100). For each query protein, the outputs of these individual predictors were used to calculate a consensus disorder profile by averaging disorder profiles of individual predictors. The outputs of these tools are represented as real numbers between 1 (ideal prediction of disorder) and 0 (ideal prediction of order). A threshold of ≥0.5 was used to identify disordered residues and regions in query proteins. A protein region was considered flexible, if its disorder propensity was in a range from 0.2 to 0.5. Results of this multiparametric computational analysis are presented in a form of mean disorder propensity calculated by averaging disorder profiles of individual predictors. Use of consensus for evaluation of intrinsic disorder is motivated by empirical observations that this approach usually increases the predictive performance compared to the use of a single predictor (96,101–108). CIDER computational platform (64) was used to generate linear net charge per residue diagrams for query proteins, whereas the sequence-based propensities of TDP-43CTD, GRN-3, and GRN-5 for granule formation associated to liquid demixing under the physiological conditions were evaluated by CatGRANULE algorithm (67), which generates positive and negative scores for proteins capable and not capable of granule formation, respectively. Finally, the presence of disorder-based binding sites in TDP-43CTD was evaluated by the ANCHOR algorithm (65,66).

## ACKNOWLEDGEMENTS

The authors would like to thank the following agencies for financial support: National Institute of Aging (1R56AG062292-01 to VR and RF1AG055088 to VNU), National Institute of General Medical Sciences (R01GM120634 to VR), and the National Science Foundation (NSF CBET 1802793 to VR). The authors also thank the National Center for Research Resources (5P20RR01647-11) and the National Institute of General Medical Sciences (8 P20 GM103476-11) from the National Institutes of Health for funding through INBRE for the use of their core facilities. The authors thank Dr. Nicholas Fawzi at Brown University for his advice on TDP-43CTD purification. The authors also thank Dr. Jonathan Lindner for his help in the use of fluorescence microscopy.

## Supplementary information

**Fig S1:**
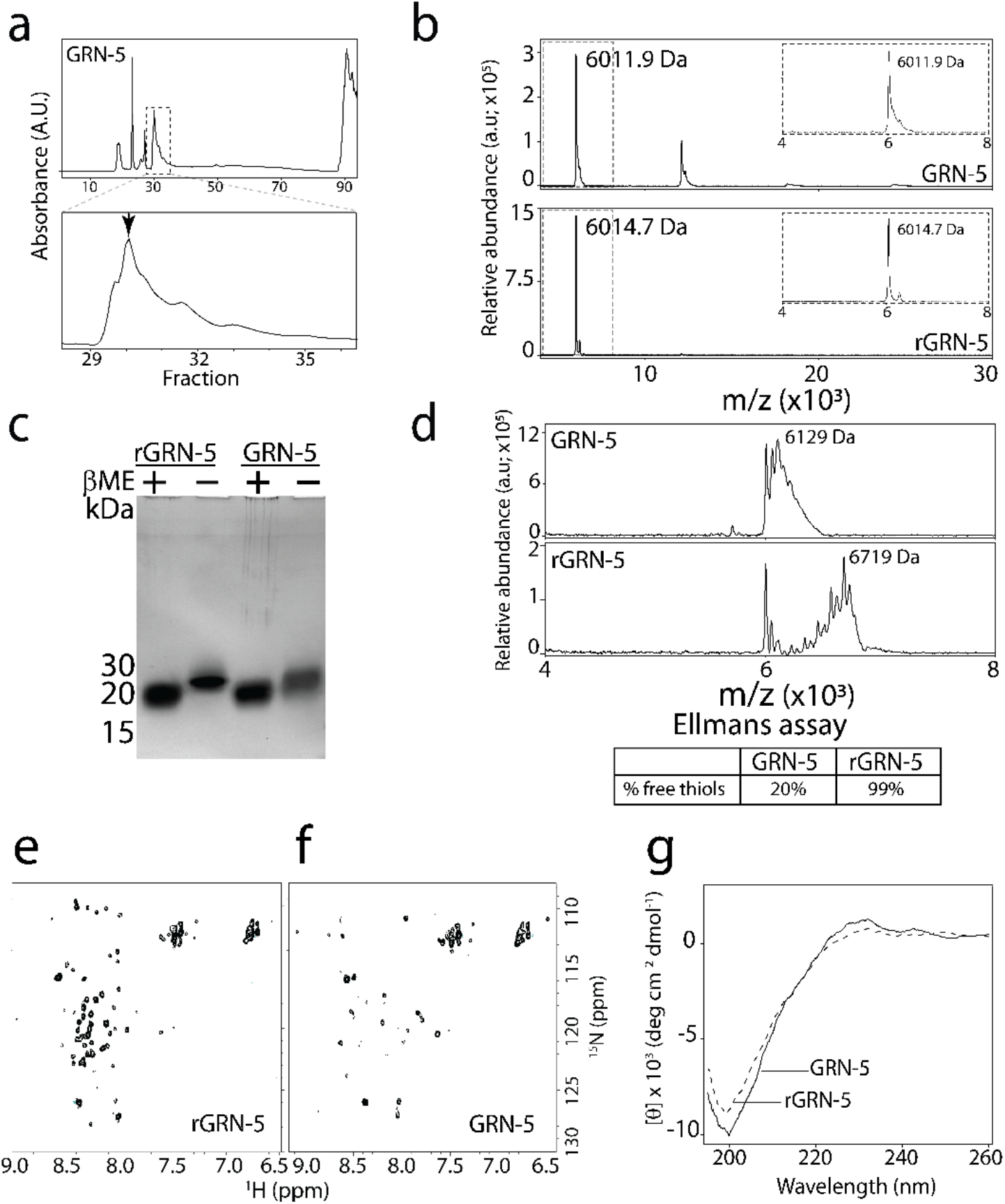
Biophysical characterization of GRN-5. a) The rHPLC profile of GRN-5 purification on a Phenomenex column. The fractions corresponding to the protein are expanded and shown below. The fraction used as the oxidized form has been highlighted with an arrow. b) The MALDI-ToF spectra of oxidized and reduced forms of GRN-5 (m.w; 6015.7 Da). c) Characterization of GRN-5 using SDS-PAGE reveals anomalous electrophoretic mobility for both forms in presence (+) and absence (-) of β-mercaptoethanol (βME). d) For determination of free thiols in respective samples, iodoacetamide labeling was performed for GRN-5 and rGRN-5. MALDI-ToF shows GRN-5 has up to two free thiols, while rGRN-5 shows 12 free cysteines. Ellman’s assay was also used where GRN-5 shows that up to 2 cysteines are free in a monomeric unit while all the 12 cysteines are present as free thiols in rGRN-5. e-f) ^15^N HMQC of the GRN-5 (e) fails to show a signal from most residues in the protein while rGRN-5 (f) shows the collapsed spectrum of a disordered protein. g) The far-UV circular dichroism spectra of the redox forms of GRN-5 show a characteristic random coil profile with a minima at 198 nm.

**Fig S2:**
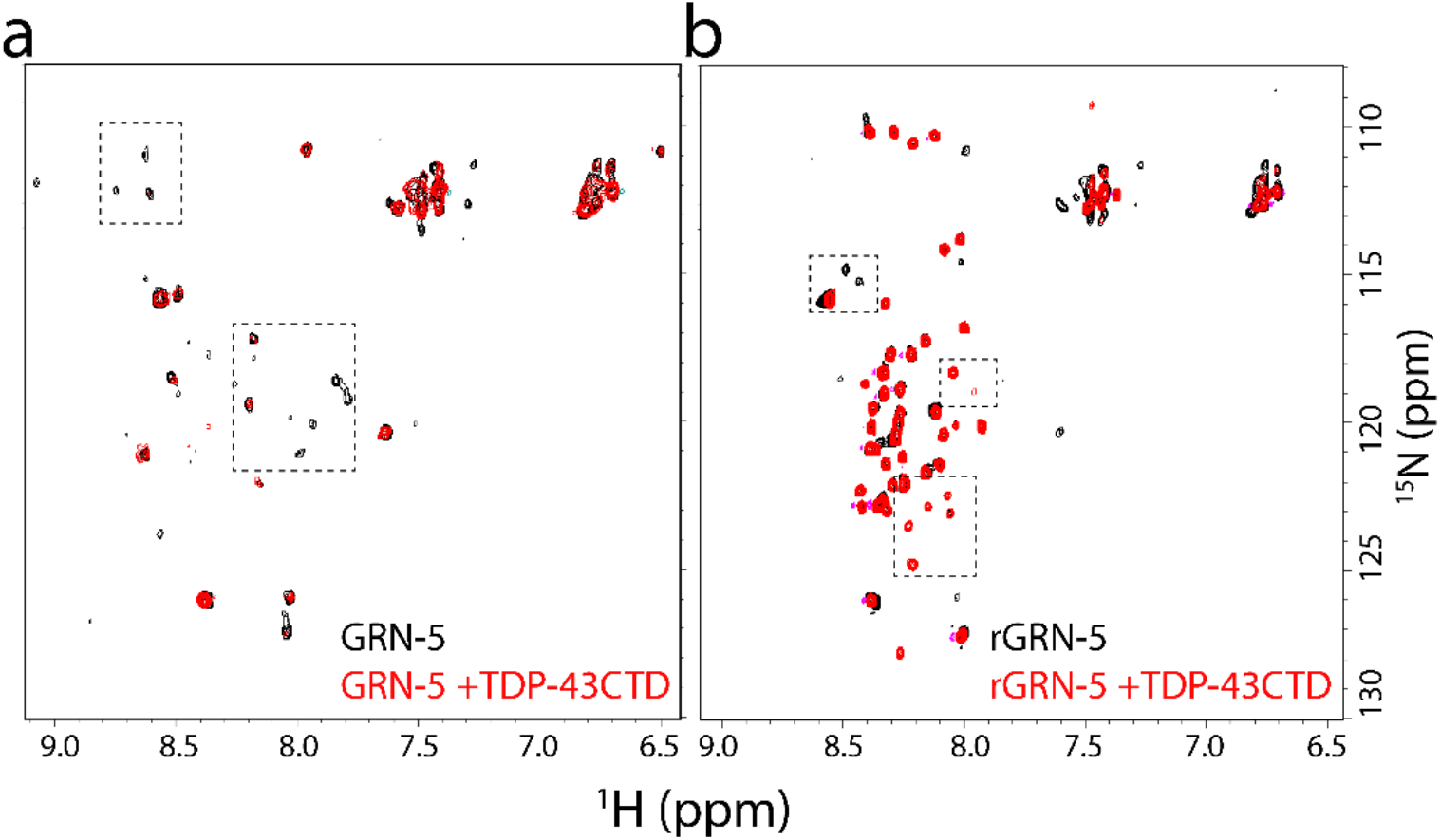
HMQC of ^15^N-GRN-5 in presence of TDP-43CTD. a-b) The HMQC spectra of 15N-GRN-5 (a) and rGRN-5 (b) in presence of TDP-43CTD shows spectral shifts for certain residues (highlighted with dashed box). The residue assignment for GRN-5 is ongoing and the exact residues involved in the interaction are not determined.

